## Supplementary material for "Revealing cancer driver genes through integrative transcriptomic and epigenomic analyses with Moonlight": S1 Fig

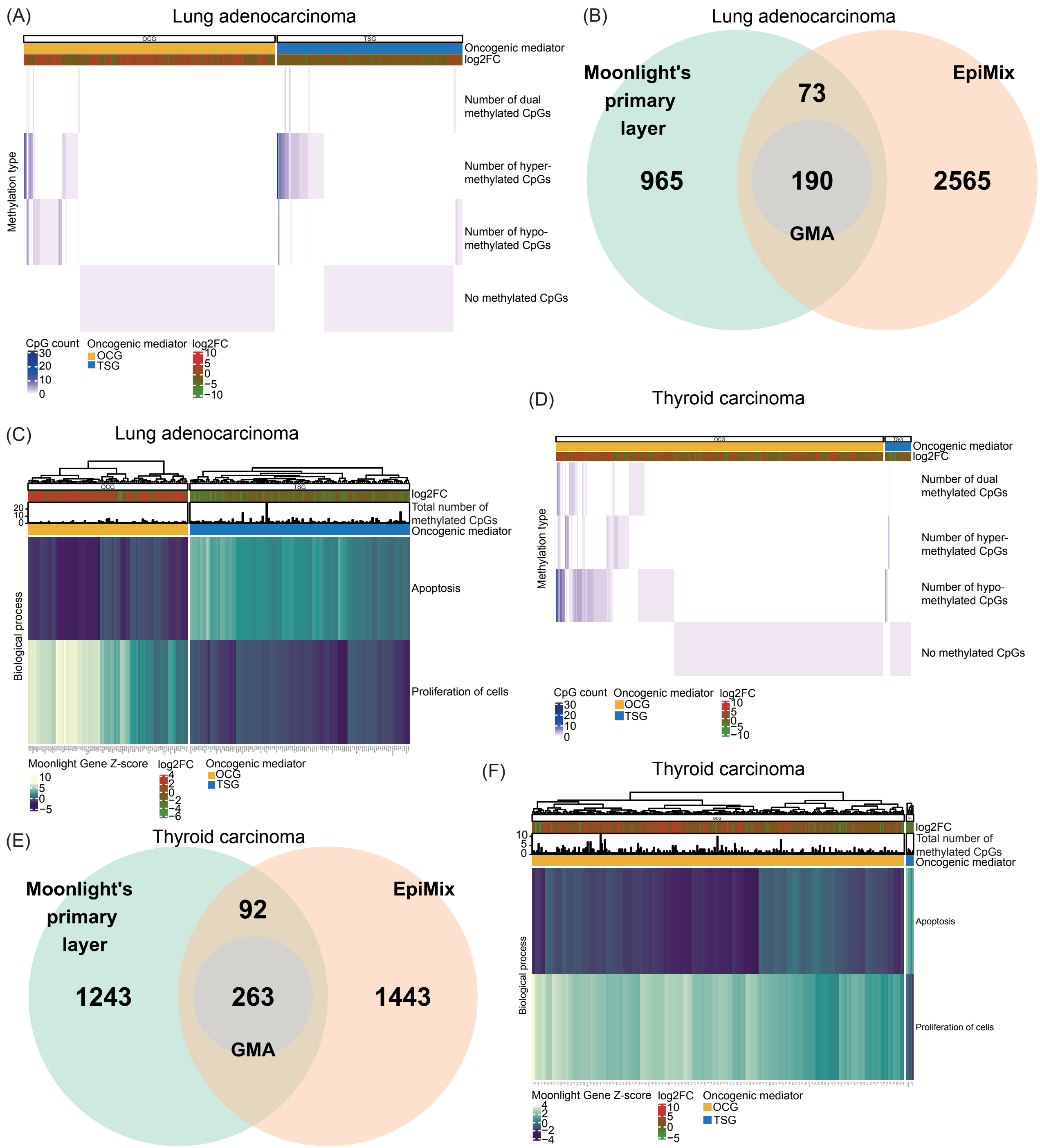


**S1 Fig. Integration of Moonlight and EpiMix for prediction of cancer driver genes. (A)** Heatmap showing number of differentially methylated CpGs and classifications of methylation status in the oncogenic mediators in lung adenocarcinoma. The heatmap was generated using the plotGMA function. **(B)** Venn diagram comparing oncogenic mediators predicted from Moonlight's primary layer with functional genes predicted from EpiMix in lung adenocarcinoma. The functional genes are genes containing differentially methylated CpG pairs whose DNA methylation state is associated with expression of the gene. Only those functional genes that contained the same methylation state in all of its associated CpGs were included in this comparison, and moreover, the dual methylation states were excluded.

**(C)** Heatmap showing the effect of the predicted driver genes in lung adenocarcinoma on apoptosis and proliferation of cells. This heatmap was generated using the function plotMoonlightMet. These effects define the basis upon which the oncogenic mediators are predicted from the PRA step in Moonlight's primary layer. **(D)** Heatmap showing number of differentially methylated CpGs and classifications of methylation status in the oncogenic mediators in thyroid carcinoma. The heatmap was generated using the plotGMA function. **(E)** Venn diagram comparing oncogenic mediators predicted from Moonlight's primary layer with functional genes predicted from EpiMix in thyroid carcinoma. The functional genes are genes containing differentially methylated CpG pairs whose DNA methylation state is associated with expression of the gene. Only those functional genes that contained the same methylation state in all of its associated CpGs were included in this comparison, and moreover, the dual methylation states were excluded. **(F)** Heatmap showing the effect of the predicted driver genes in thyroid carcinoma on apoptosis and proliferation of cells. This heatmap was generated using the function plotMoonlightMet. These effects define the basis upon which the oncogenic mediators are predicted from the PRA step in Moonlight's primary layer.
