## Supplementary material for "Revealing cancer driver genes through integrative transcriptomic and epigenomic analyses with Moonlight": S2 Fig

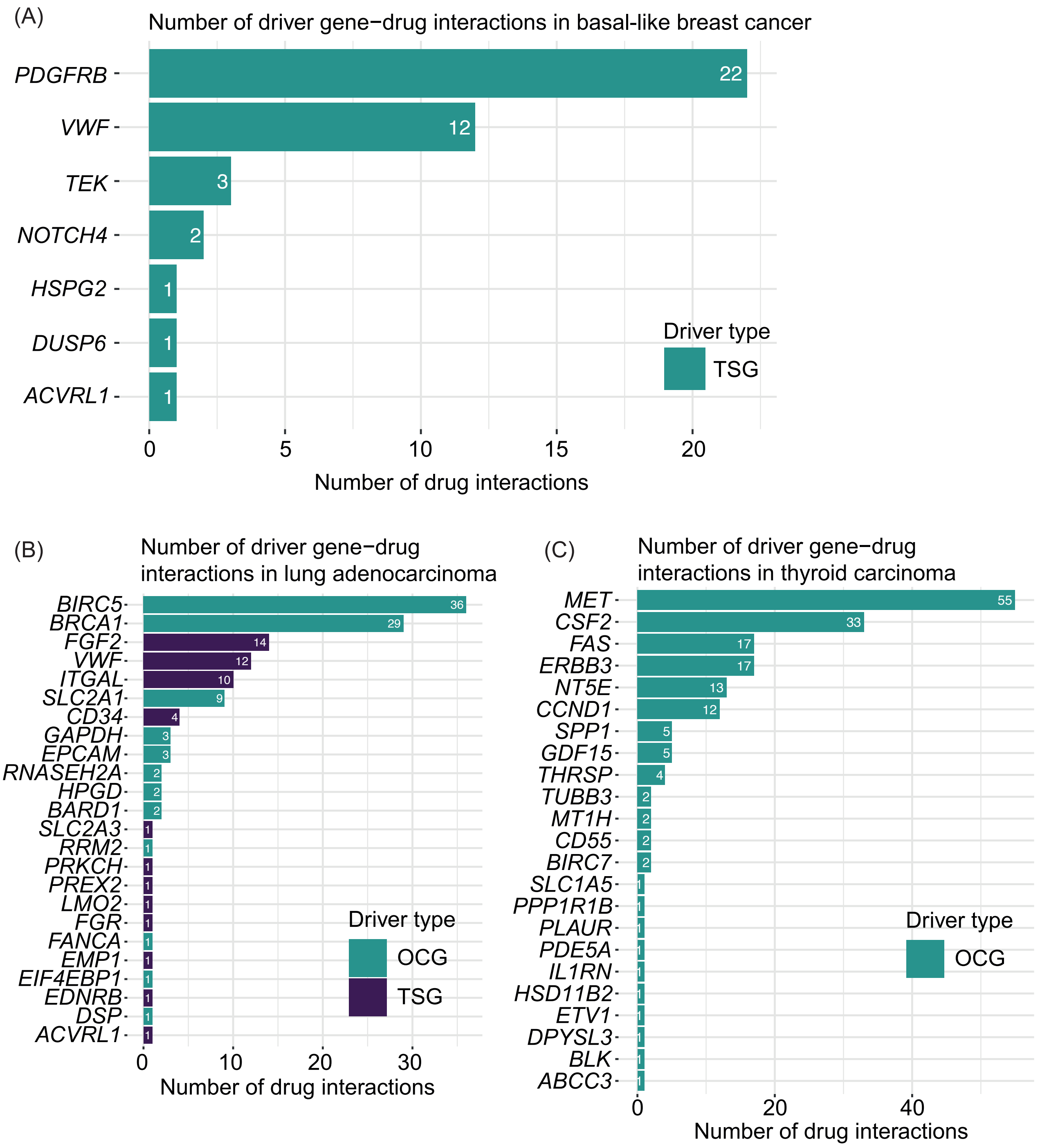


**S2 Fig. Number of driver gene-drug interactions.** Number of driver gene-drug interactions in **(A)** basal-like breast cancer, **(B)** lung adenocarcinoma, and **(C)** thyroid carcinoma found by querying DGIdb. The driver genes are stratified into OCGs and TSGs. The number of drug interactions is shown on the *x* axis and the driver genes are shown on the *y* axis.
