## Supplementary material for "Revealing cancer driver genes through integrative transcriptomic and epigenomic analyses with Moonlight": S1 Text

### Evidence integration for defining driver genes

Gene Methylation Analysis (GMA) predicts driver genes by comparing EpiMix's predictions of methylation state and Moonlight's predictions of driver role. This is based on the assumption with which methylation changes can activate or inactivate driver genes. Suppose EpiMix predicts a gene to contain one or more hypermethylated CpG sites whereas Moonlight's primary layer predicts this gene to be an oncogene. This indicates conflicting evidence between the mechanism of (in)activation and putative driver role between the two tools. Consequently, this gene is labeled with "conflicting" evidence. Contrary, if EpiMix for instance predicts a gene to be associated with hypermethylated CpG site(s) and Moonlight predicts this gene as a tumor suppressor, this gene is labeled with an "agreement" evidence to signify correspondence between EpiMix and Moonlight. Those oncogenic mediators labeled with an "agreement" evidence are retained as the final set of predicted driver genes.
