## Supplementary material for "Revealing cancer driver genes through integrative transcriptomic and epigenomic analyses with Moonlight": S2 Text

### Methods of case study: Prediction of driver genes with differential methylation in basal-like breast cancer, lung adenocarcinoma, and thyroid carcinoma using Moonlight

#### Application of new functionality to three cancer (sub)types

To illustrate the new functionality in Moonlight, Gene Methylation Analysis (GMA), and predict driver genes driven by methylation changes, we applied Moonlight on three cancer (sub)types: basal-like breast cancer, lung adenocarcinoma, and thyroid carcinoma. We obtained gene expression, methylation, and clinical data from The Cancer Genome Atlas (TCGA) using the R Bioconductor package TCGAbiolinks [1-3]. For basal-like breast cancer, we retrieved this particular subtype using the PanCancerAltas_subtype function in TCGAbiolinks. We processed the gene expression data as previously described using TCGAbiolinks [1-5] and the methylation data using the Preprocess_DNAMethylation function from EpiMix as previously done [6-7]. We kept only patients who had both expression and methylation data resulting in 144, 474, and 551 patients for basal-like breast cancer, lung adenocarcinoma, and thyroid carcinoma, respectively. Subsequently, we performed differential expression analysis (DEA) between each of the cancer (sub)types and corresponding normal samples using limma-voom including tissue source site (TSS) as a covariate in the linear modelling to account for TSS as a batch factor as previously outlined [4-5]. Finally, we applied the Moonlight pipeline including the new GMA functionality and predicted driver genes using apoptosis and cell proliferation as the underlying biological foundation given the recognized role of these two processes in carcinogenesis [4-5].

#### Enrichment analysis of predicted driver genes

We performed enrichment analyses of predicted driver genes using the enrichr R package [8-10] and the "MSigDB Hallmark 2020" database. We used the enrichr function to perform enrichment analyses on the predicted driver gene sets of each cancer (sub)type and selected significantly enriched terms as those with an adjusted $p$-value < 0.05.

#### Survival analysis of predicted driver genes

We conducted survival analysis of predicted driver genes in the three cancer (sub)types using Cox proportional hazards regression and Kaplan-Meier survival analysis implemented in the survival [11-12], survminer [13], and survMisc [14] R packages. We used the last follow-up or death time and vital status (dead or alive) of the patients as the survival data. First, we tested the proportional hazards assumption using the cox.zph function and proceeded with genes satisfying this assumption. For these genes, we next fit a univariate Cox regression model using the coxph function with survival data as the response variable and expression values of the respective gene as a continuous explanatory variable. Following FDR correction of resulting *p*-values, genes whose expression had a significant effect on survival were retained (FDR < 0.05) and subject to subsequent multivariate Cox regression analysis. Here, tumor stage, age of patients, and sex of patients were included as additional explanatory variables to ensure that the effect of the gene on survival was in fact due to its expression and not because of confounding factors such as stage, age, and sex. Sex of patients were not included for basal-like breast cancer as all patients were female. Genes whose expression showed a significant effect on survival at the multivariate level (*p*-value < 0.05) were considered prognostic. Moreover, we performed Kaplan-Meier survival analysis on these prognostic genes to also evaluate differences in survival between two discrete expression groups. Patients with expression values above and below the median expression level of the respective gene were divided into a high and low expression group, respectively. We fit survival curves using the discrete expression group as the explanatory variable and evaluated the significance of difference in survival between the two groups by means of a log-rank test as *p*-value < 0.05.

**Exploration of predicted driver genes as drug targets**

To investigate driver gene-drug interactions, we applied the R package rDGIdb [15-16] to query the Drug-Gene Interaction Database (DGIdb) 5.0 [17] using only cancer-specific data sources: Cancer Genome Interpreter (CGI) [18], Clinical Interpretation of Variants in Cancer (CIViC) [19], Catalogue of Somatic Mutations in Cancer (COSMIC) [20], CancerCommons, ClearityFoundationBiomarkers, ClearityFoundationClinicalTrial, Database of Curated Mutations (DoCM) [21], JAX Clinical Knowledgebase (JAX-CKB) [22], MyCancerGenome [23], MyCancerGenomeClinicalTrial, National Cancer Institute (NCI), Oncology Knowledge Base (OncoKB) [24], and TALC.

**Comparison with Driver Mutation Analysis (DMA), previously available secondary layer of evidence for Moonlight**

We previously developed a different secondary layer of evidence for Moonlight called Driver Mutation Analysis (DMA) [5] which provides complementary evidence with respect to GMA by using a different mechanistic indicator. DMA uses cancer-associated mutation data available for the predicted oncogenic mediators to assess their driver status, distinguishing between driver and passenger mutations by means of the CScape-somatic predictor [25]. DMA needs to be used after the Pattern Recognition Analysis (PRA) step in the Moonlight workflow and requires 1) the set of differentially expressed genes (DEGs) used in the primary layer, 2) the predicted oncogenic mediators from Moonlight's primary layer, and 3) a mutation annotation file (MAF) which contains all the mutations of interest. Here, our mutation data only included mutations from patient samples used in the Moonlight primary layer analysis and GMA, for consistency. We performed the DMA with default options using the aforementioned inputs. To compare the predictive capabilities of DMA and GMA in identifying driver genes, we utilized the R package VennDiagram [26] assessing the overlap between the sets of driver genes predicted by each method (DMA or GMA). Additionally, we performed enrichment analysis of driver genes predicted by DMA similarly as we did for GMA. We compared the significantly enriched terms from DMA to the ones identified by GMA to understand the overlap in the cancer hallmarks that the two methods identified.

### References


1. Colaprico A, Silva TC, Olsen C, et al. TCGAbiolinks: An R/Bioconductor package for integrative analysis of TCGA data. Nucleic Acids Res 2016; 44:1–11
2. Mounir M, Lucchetta M, Silva TC, et al. New functionalities in the TCGAbiolinks package for the study and integration of cancer data from GDC and GTEX. PLoS Comput Biol 2019; 15:1–18
3. Silva TC, Colaprico A, Olsen C, et al. TCGA Workflow: Analyze cancer genomics and epigenomics data using Bioconductor packages. F1000Res 2016; 5:1–61
4. Colaprico A, Olsen C, Bailey MH, et al. Interpreting pathways to discover cancer driver genes with Moonlight. Nat Commun 2020; 11:1–17
5. Nourbakhsh M, Saksager A, Tom N, et al. A workflow to study mechanistic indicators for driver gene prediction with Moonlight. Brief Bioinform 2023; 24:1–13
6. Zheng Y, Jun J, Brennan K, et al. EpiMix is an integrative tool for epigenomic subtyping using DNA methylation. Cell Reports Methods 2023; 3:1–23
7. Gevaert O, Tibshirani R, Plevritis SK. Pancancer analysis of DNA methylation-driven genes using MethylMix. Genome Biol 2015; 16:1–13
8. Chen EY, Tan CM, Kou Y, et al. Enrichr: interactive and collaborative HTML5 gene list enrichment analysis tool. BMC Bioinformatics 2013; 14:1–14
9. Xie Z, Bailey A, Kuleshov M V., et al. Gene Set Knowledge Discovery with Enrichr. Curr Protoc 2021; 1:1–84
10. Kuleshov M V., Jones MR, Rouillard AD, et al. Enrichr: a comprehensive gene set enrichment analysis web server 2016 update. Nucleic Acids Res 2016; 44:W90–W97
11. Therneau T. A package for survival analysis in R. R package, https://CRAN.R-project.org/package=survival 2022
12. Therneau TM, Grambsch PM. Modeling Survival Data: Extending the Cox Model. 2000
13. Kassambara A, Kosinski M, Biecek P, et al. survminer: Drawing Survival Curves using 'ggplot2'. R package, https://CRAN.R-project.org/package=survminer 2021
14. Dardis C. survMisc: Miscellaneous Functions for Survival Data. R package, https://CRAN.R-project.org/package=survMisc 2022

15. Thurnherr T, Singer F, Stekhoven DJ, et al. Genomic variant annotation workflow for clinical applications. F1000Res 2016; 5:1–13
16. Wagner AH, Coffman AC, Ainscough BJ, et al. DGIdb 2.0: Mining clinically relevant drug-gene interactions. Nucleic Acids Res 2016; 44:D1036–D1044
17. Cannon M, Stevenson J, Stahl K, et al. DGIdb 5.0: rebuilding the drug-gene interaction database for precision medicine and drug discovery platforms. Nucleic Acids Res 2024; 52:D1227–D1235
18. Tamborero D, Rubio-Perez C, Deu-Pons J, et al. Cancer Genome Interpreter annotates the biological and clinical relevance of tumor alterations. Genome Med 2018; 10:25
19. Griffith M, Spies N, Krysiak K, et al. CIViC is a community knowledgebase for expert crowdsourcing the clinical interpretation of variants in cancer. Nat Genet 2017; 47:7
20. Tate JG, Bamford S, Jubb HC, et al. COSMIC: the Catalogue Of Somatic Mutations In Cancer. Nucleic Acids Res 2019; 47:D1
21. Ainscough B, Griffith M, Coffman A, et al. DoCM: a database of curated mutations in cancer. Nat Methods 2016; 13:10
22. Patterson SE, Liu R, Statz CM, et al. The clinical trial landscape in oncology and connectivity of somatic mutational profiles to targeted therapies. Hum Genomics 2016; 10:4
23. Yeh P, Chen H, Andrews J, et al. DNA-Mutation Inventory to Refine and Enhance Cancer Treatment (DIRECT): a catalog of clinically relevant cancer mutations to enable genome-directed anticancer therapy. Clin Cancer Res 2013; 19:7
24. Chakravarty D, Gao J, Phillips SM, et al. OncoKB: A Precision Oncology Knowledge Base. JCO Precis Oncol 2017; 1
25. Rogers MF, Gaunt TR, Campbell C. CScape-somatic: distinguishing driver and passenger point mutations in the cancer genome. Bioinformatics 2020; 36:12
26. Chen H, Boutros PC. VennDiagram: a package for the generation of highly-customizable Venn and Euler diagrams in R. BMC Bioinformatics 2011; 12:35
